# Design and Validation of a 3D-Printed Motorized Biaxial Cell Stretching Device

**DOI:** 10.64898/2026.06.25.734357

**Authors:** Nada M. Kafour, Noor A. Al-Maslamani, Badria F. Al-Sammak, Henning F. Horn

## Abstract

Mechanical forces have a major effect on cell behavior. Most cells in vitro are grown under static conditions on hard tissue culture plastic, conditions that do not accurately reflect living tissues. The ability of cells to sense and respond to mechanical forces is essential for key biological processes, including development, proliferation, and migration. Disruption of the ability to respond to mechanical forces are known to be a critical factor in many diseases, including cardiovascular disease, progeria, and cancer. Here, we present the design, fabrication, and biological testing of a custom-built cell-stretching device that applies controlled biaxial strain to cells cultured on a polydimethylsiloxane (PDMS) membrane. We then used this device to examine how cells respond to strain. In response to biaxial strain, MCF-7 cells activated the mechanosensitive immediate early gene (IEX-1), with its expression increasing significantly after 1 and 3 hours of stretching. Cells exposed to mechanical strain also remodeled their cytoskeleton in a direction-dependent manner. Under uniaxial strain, actin filaments reoriented perpendicular to the stretch direction, whereas biaxially stretched cells do not promote directional reorientation, but instead appear to reinforce actin at the cell periphery. Similarly, cells under uniaxial strain exhibited changes in nuclear orientation and shape that were not observed under biaxial strain. Nuclear area remained unchanged in either strain condition. These results highlight that the biaxial stretcher can be used to apply strain to cells, and that cells respond differently to biaxial strain compared to what has been reported for uniaxial strain.

## Introduction

Most cells in the human body experience mechanical forces and have the ability to sense and respond to mechanical stimuli (Maurer & Lammerding, 2019a). The process by which cells translate mechanical forces into signaling events is known as mechanotransduction (Martino et al., 2018) and can influence proliferation, differentiation, migration, and apoptosis (Janota et al., 2020). As such, mechanical strain shapes the physiology of most organs in the human body.

One major route of mechanotransduction begins with a mechanical stimulus in the cellular environment that is transmitted into the nucleus through a series of intermediary signaling components. For example, mechanical forces acting on the extracellular matrix (ECM) are sensed by integrins and focal adhesions and transmitted through the interconnected network of cytoskeletal elements, including actin filaments, microtubules, and intermediate filaments (Martino et al., 2018). From the cytoskeleton, mechanical force is then transmitted across the nuclear envelope through the LINC complex to the nuclear lamina and chromatin (Hieda, 2019; Lityagina & Dobreva, 2021; Maurer & Lammerding, 2019a). This force transmission can alter nuclear shape, chromatin organization and accessibility, and gene expression (Lityagina & Dobreva, 2021; Maurer & Lammerding, 2019b; Uhler & Shivashankar, 2017).

Altered cellular responses to mechanical forces are key contributors to many diseases, including laminopathies, cardiovascular diseases, and cancer (Di et al., 2023; Donnaloja et al., 2020; Lityagina & Dobreva, 2021; Maurer & Lammerding, 2019a; Yamashiro & Yanagisawa, 2020). Indeed, cells in which mechanotransduction is impaired have contributed to our understanding of disease mechanisms and highlight potential therapeutic approaches (Di et al., 2023).

Experimental techniques that can mimic *in vivo* mechanical strain have been key in elucidating cellular mechanotransduction pathways (Muhamed et al., 2017). A common technique uses cell stretching systems, which exert varying levels of mechanical strain to model different mechanotransduction responses. These systems work by deforming a compliant culture substrate and impose uniaxial or biaxial strain on a 2D monolayer of cells (Brown, 2000; Constantinou & Bastounis, 2023; Kamble et al., 2016a). Strain magnitude in these systems is often expressed as a percentage change in substrate length along the strain axis (Kamble et al., 2016a). Physiological strain levels vary widely among tissues. Bone usually experiences relatively low levels of strain, whereas tissues such as the myocardium, lung parenchyma, and blood vessels can experience larger cyclic deformations under normal conditions (Kamble et al., 2016b; Tschumperlin & Margulies, 1998).

Uniaxial stretching applies force along a single axis and has been a widely used system for mechanobiology studies (Al-Maslamani et al., 2021; Atcha et al., 2018; Fang et al., 2022; Kah et al., 2021; Kamble et al., 2017; Kreutzer et al., 2020; Shao et al., 2013; Yadav et al., 2023; Yost et al., 2000). Several commercial uniaxial stretchers are available and include STREX and Flexcell systems. In addition, a large number of custom designs have been reported, including LEGO-based and 3D-printed motorized stretchers (Al-Maslamani et al., 2021; Boulter et al., 2020). These devices are especially relevant for cell types and tissues that normally experience mostly uni-directional tension, such as striated muscle cells, tendons, and ligaments (Salameh et al., 2010; Zitnay & Weiss, 2018). However, the mechanical environment experienced by most cells *in vivo* is rarely uniaxial. Many tissues, including the heart, blood vessel walls, lung epithelium, and skin, experience strain in multiple directions (Brady et al., 2022; Friedrich et al., 2019; Kassab, 2006; Nagle et al., 2023; Roan & Waters, 2011). In order to more closely mimic these strain conditions, biaxial stretching devices have been developed.

Biaxial stretchers deform the substrate along two orthogonal axes, either equally (equibiaxial) or unequally (non-equibiaxial), to better represent multidirectional forces (Constantinou & Bastounis, 2023). Commercial systems are available from STREX and Flexcell biaxial systems. While these systems are well-established, they are often expensive and difficult to customize for specific experiments. Custom biaxial stretchers can reduce cost and improve design flexibility. Several systems use vacuum or indenters to pull membranes across a stationary loading post, thereby applying equibiaxial strain (Imsirovic et al., 2015; Ursekar et al., 2014). These systems do not generally allow for variations between axes. In addition, strain can vary across the membrane, depending on which part of the membrane is examined (Imsirovic et al., 2015; Rana et al., 2008; Ursekar et al., 2014). Other custom biaxial stretching systems have small culture areas that limit the number of cells available for downstream analysis (Tremblay et al., 2013; Win et al., 2017). We therefore sought to develop a biaxial stretcher that provides a more uniform strain across a larger culture area, suitable for both imaging and biochemical analysis.

We previously developed a 3D-printed, motorized uniaxial stretching system for applying controlled mechanical strain (Al-Maslamani et al., 2021). Building on this platform, we sought to develop a cost-effective biaxial stretcher that retains the same PDMS membrane and motor type but could stretch independently along two perpendicular axes. Here we report its design, fabrication, mechanical validation, and assess its compatibility with cell culture, microscopy, and biochemical analysis. We also report a direct comparison of cellular responses between uniaxial and biaxial strain using the two systems developed in our group.

## Results

### Design, assembly, and control of the biaxial cell stretcher

The biaxial stretcher was developed to apply strain to adherent cells cultured on PDMS membranes (Figure 1). All structural components were designed in Autodesk Inventor (Figure 1A), exported as STL files, and printed on an Ultimaker 3D printer. The assembled device contains four micro linear motors arranged around a square PDMS membrane (Figure 1B). Each motor arm connects to a motor hand attached to one corner of the membrane. During biaxial stretching, opposing motor pairs move outward to deform the membrane along the X and Y axes. Because each motor is connected independently, the system could potentially also be programmed for non-equibiaxial loading, though we do not describe this here.

**Figure 1.**
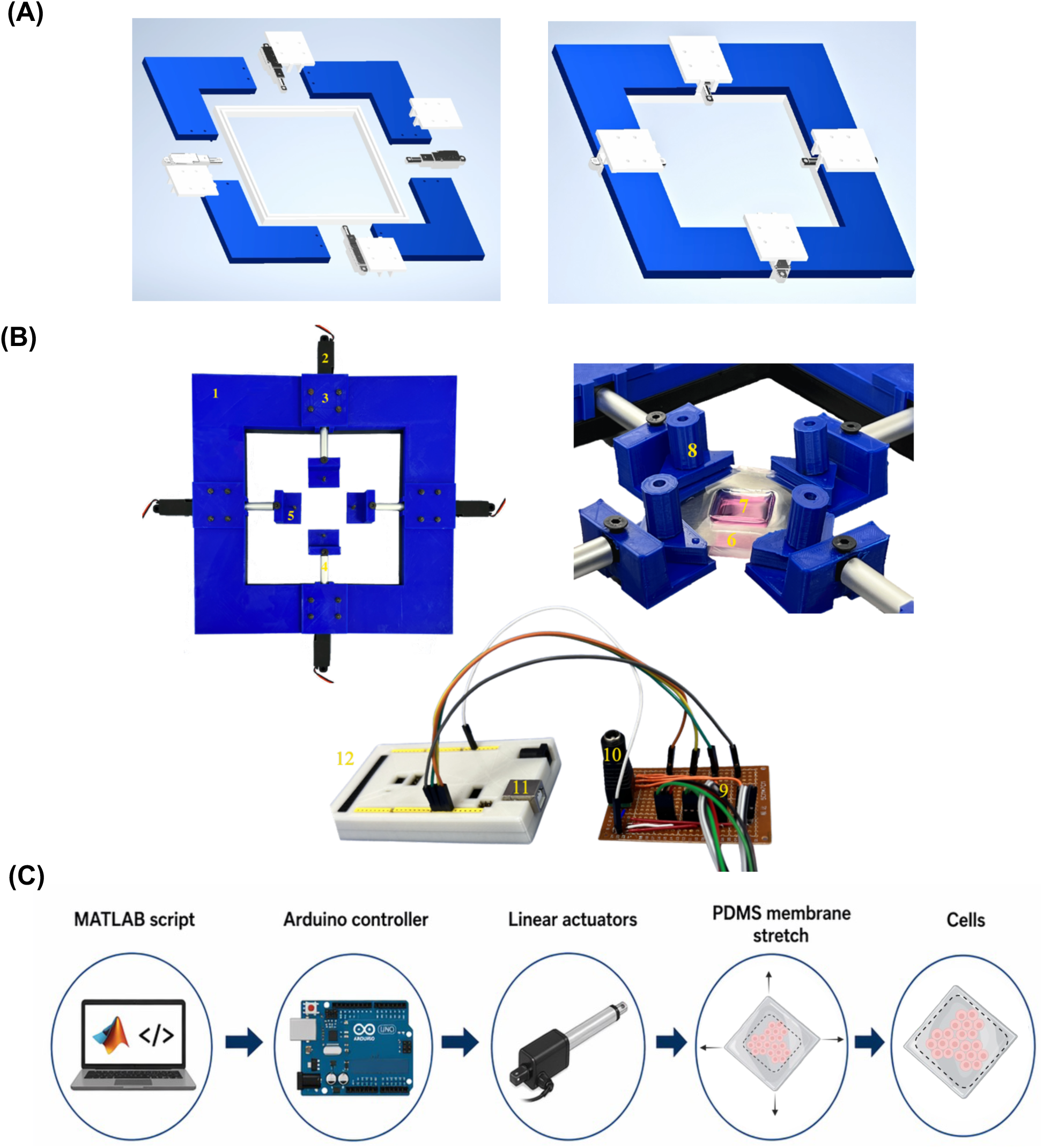
Design and workflow of the biaxial cell stretching device. **(A)** Computer-aided design (CAD) model of the biaxial cell stretcher base designed in Autodesk Inventor. **(B)** Assembled biaxial stretcher showing mechanical and electrical components: (1) 3D-printed PLA stretcher base; (2) micro linear motor; (3) motor cover; (4) motor arm; (5) motor hand, for PDMS membrane attachment; (6) PDMS membrane mounted onto the biaxial stretcher displaying the membrane well; (7) Fibronectin-coated culture well containing cells; (8) triangular clamps and screws; (9) circuit board; (10) electrical power supply; (11) USB port; (12) Arduino Mega 2560. **(C)** Schematic of the biaxial stretcher control workflow.

The PDMS membrane contains a central culture well (coated with fibronectin for cell attachment) and is secured to the motor hands using screws and triangular clamps. This attachment method evenly distributes clamping force at the membrane corners in order to prevent membrane tearing during repeated stretching cycles

The four motors are connected to a shared circuit board and powered by an external power supply. Motor movement is controlled using an Arduino Mega 2560 connected to a computer by USB. The control workflow was adapted from Al-Maslamani et al. (2021). The original MATLAB code for the uniaxial stretcher was modified to control four motors through digital pins D8, D9, D10, and D11. The complete uniaxial and biaxial MATLAB code is available at https://github.com/Henning-Horn.

Stretching parameters are entered in MATLAB and include motor displacement, experiment duration, stretch mode, and hold time. In static mode, the motors move the membrane to a target position and hold it there. In cyclic mode, the motors repeatedly move the membrane between relaxed and stretched positions. In this study, the motor speed is 6.5 mm/s and cyclic stretch was applied at 0.5 Hz, corresponding to one complete cycle every 2 seconds.

When choosing the strain percentage in our experiments we chose 7% for biaxial stretching. Equibiaxial strain has been shown to induce biochemical responses at magnitudes as low as 1-5%, and studies using 5-10% cyclic strain have reported changes in cytoskeletal organization, cell spreading, and nuclear morphology (Cui et al., 2015; Geiger et al., 2006; Roshanzadeh et al., 2020). Therefore, 7% was selected as the strain magnitude based on the literature. In addition, we found that strains above 10% caused rupture of the PDMS membrane. The selected strain remained well below the membrane’s mechanical limit, reducing the risk of failure during cyclic stretching, while promoting a strain application with predicted measurable cellular changes.

### Fabrication and mechanical validation of the PDMS membrane

The choice of membrane material is a key design aspect in a cell-stretching device, as it affects strain uniformity, cell attachment, and overall biocompatibility (Constantinou & Bastounis, 2023; Kamble et al., 2016c). PDMS is the most widely used substrate in custom-built stretching systems. It is easy to mold into different shapes and is suitable for microscopy due to its transparency. The PDMS membranes we used were fabricated using a modified version of the protocol described by Al-Maslamani et al. (2021). We modified the casting method in order to minimize the variation in well thickness. We used a commercially prepared PDMS sheet with a thickness of 250 µm, placed at the base of the mold, and uncured PDMS was then poured over the sheet to form the central well structure (Figure 2A). The PDMS membrane mold was designed for use with both the uniaxial and biaxial stretchers. To adapt the membrane for the biaxial device, the cured PDMS was cut into a square, and a hole was created at each corner for attachment to the four motor arms (Figure 2A).

**Figure 2.**
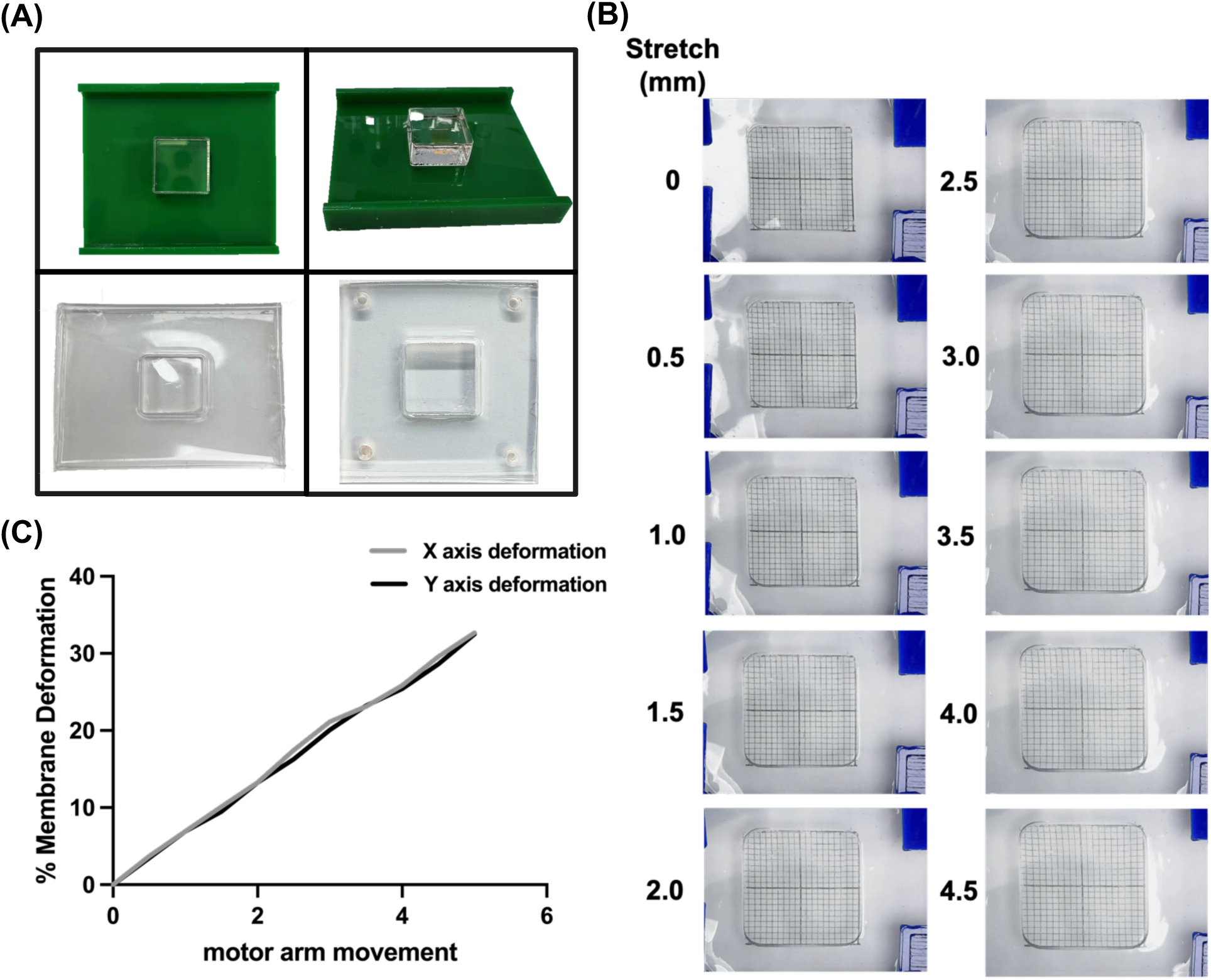
Fabrication and deformation analysis of the PDMS membrane. **(A)** Images of the Plexiglass mold used to cast the PDMS membrane and images of the cured PDMS membrane. The bottom right PDMS membrane has been cut and hole-punched for biaxial stretching. **(B)** Representative images of the PDMS membrane during progressive biaxial stretching as motor arm movement increased from 0 to 4.5 mm. **(C)** Quantification of membrane deformation along the X and Y axes in response to increasing motor arm movement. Percent membrane deformation is plotted against motor arm movement, showing similar deformation along both axes.

To validate the mechanical performance of the membranes, we characterized deformation during biaxial stretching. Membrane deformation along the X and Y axes was quantified by measuring displacement across each axis at increasing motor arm movements (Figure 2B). The biaxial stretcher PDMS membrane started at 0mm motor arm movement, and the x- and y-axes were measured at 18.39 mm and 18.43 mm, respectively, then increased by 0.5 mm increments. At each stretch interval, the membrane was measured, and the strain distribution across its surface was assessed using a grid pattern transferred onto the membrane before stretching. The membrane deformation graph showed almost equal deformation in both the x and y axes across all stretch increments (Figure 2C).

### Biaxial stretching induces time-dependent upregulation of IEX-1 in MCF-7 cells

To determine whether stretching in the biaxial stretcher altered gene expression, we measured expression of IEX-1 following cyclic biaxial strain application. MCF-7 cells on fibronectin-coated PDMS membranes were left unstretched or subjected to 7% (1mm motor arm movement) strain for 30 minutes, 1 hour, or 3 hours, and IEX-1 expression was quantified by real-time PCR relative to PGK-1 and HPRT (Figure 3). IEX-1 expression showed no significant change after 30 minutes of stretching (Figure 3A). Expression then increased significantly, rising approximately 1.5-fold relative to the unstretched control at 1 hour and approximately 2.3-fold at 3 hours (Figure 3B and C). This upregulation indicates that cells on the PDMS membrane were able to sense the applied biaxial strain and responded with a transcriptional response.

**Figure 3.**
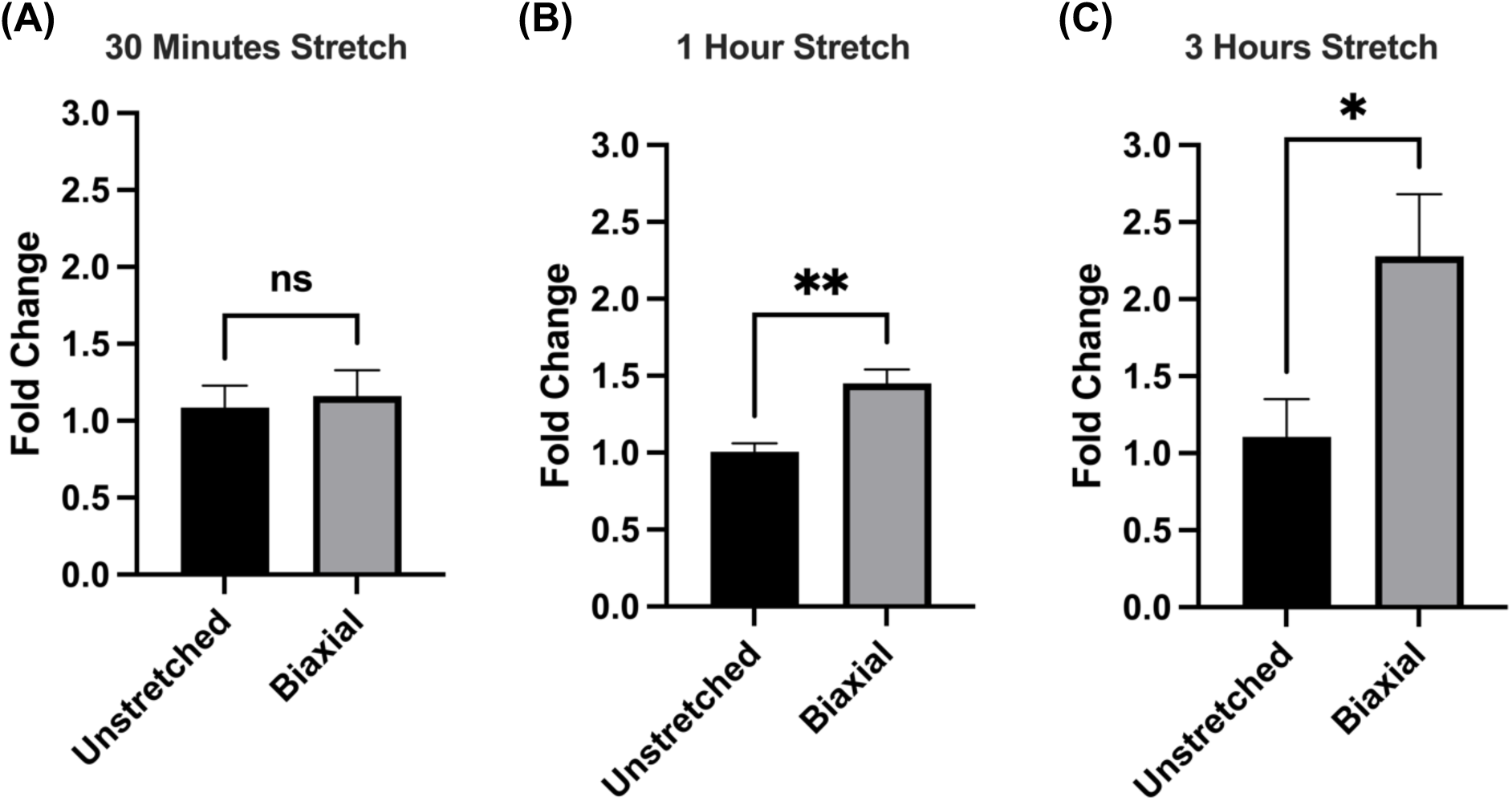
Real-time PCR analysis of IEX-1 in biaxially stretched cells. IEX-1 expression levels in unstretched cells and cells biaxially stretched at 1mm motor arm movement for **(A)** 30 minutes, **(B)** 1 hour, and **(C)** 3 hours. IEX-1 expression was quantified by real-time PCR and normalized to the housekeeping genes PGK-1 and HPRT. Data are presented as mean ± SEM (n = 6). Statistical significance was assessed using Welch’s t-test.

### Actin cytoskeletal remodeling in response to uniaxial and biaxial stretch

Uniaxial cyclic stretch has been shown to reorganize actin stress fibers and alter cell orientation (Hayakawa et al., 2001; Tondon & Kaunas, 2014; Wang et al., 2001). To compare actin organization under different stretch patterns, MCF-7 cells were left unstretched or subjected to uniaxial or biaxial cyclic stretch for 6 hours. Cells were fixed immediately after stretching and stained with phalloidin and DAPI to visualize F-actin and nuclei, respectively (Figure 4A). Unstretched cells showed dispersed actin staining throughout the cytoplasm. After uniaxial strain, actin fibers appeared more aligned, with stronger orientation perpendicular to the strain direction. In biaxially stretched cells, actin staining appeared more pronounced near the cell periphery and around the nucleus, but without the same directional alignment observed after uniaxial strain (Figure 4A). Images were analyzed in ImageJ using the directionality tool and this quantitative analysis supported the visual observations: uniaxially stretched cells showed a stronger peak at 90°, whereas unstretched and biaxially stretched cells showed roughly equal distribution across all angles (Figure 4B).

**Figure 4.**
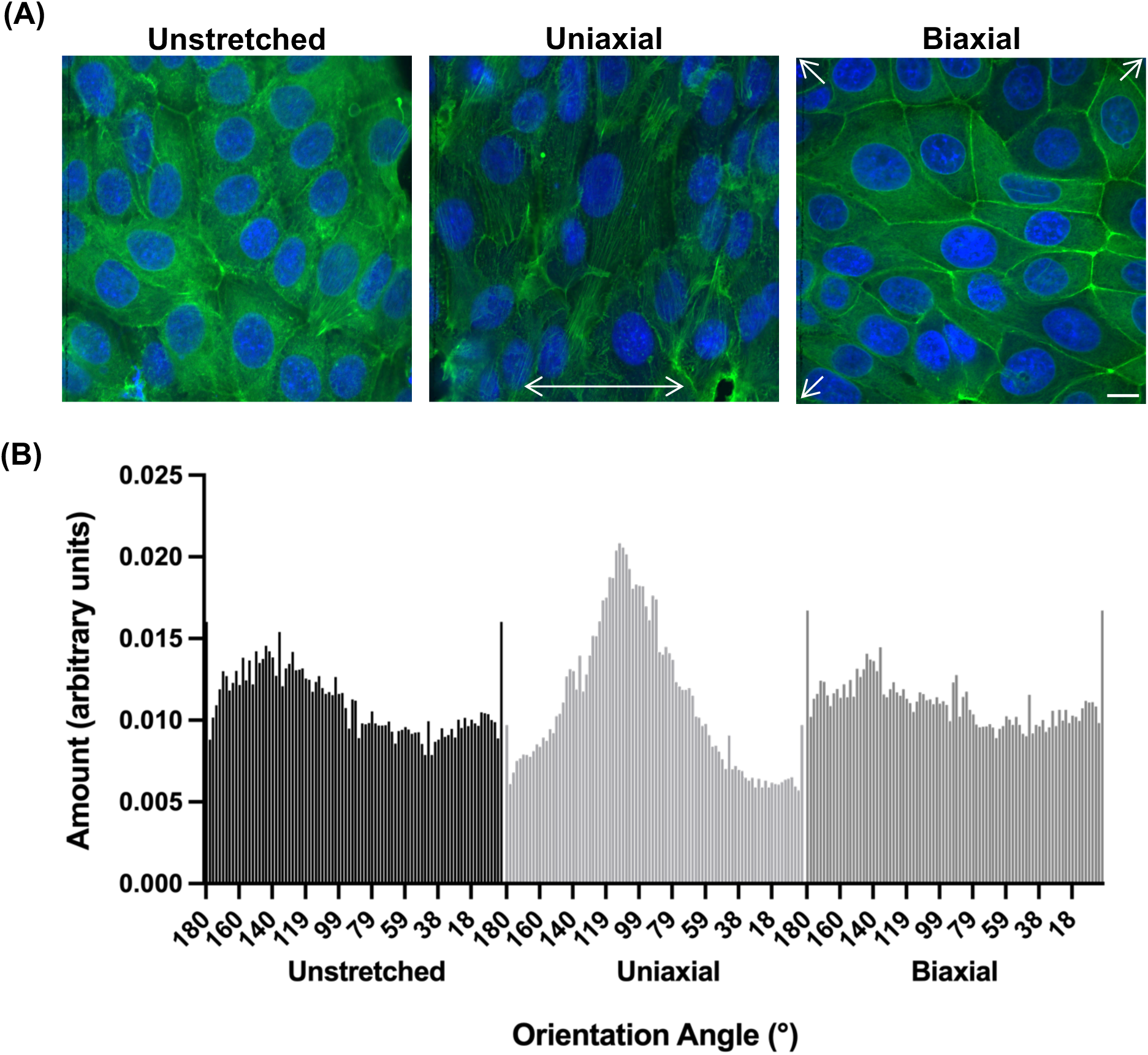
Actin cytoskeletal organization in MCF-7 cells after mechanical stretching. **(A)** Representative 60x confocal images of MCF-7 cells maintained under unstretched (control), uniaxial, and biaxial stretched conditions for 6 hours. Following fixation, F-actin and nuclei were visualized by staining with phalloidin (green) and DAPI (blue). Scale bar represents 20 μm **(B)** Histogram displaying actin filament orientation angle distribution after cyclic stretching. Directionality is expressed relative to the uniaxial stretch axis (0° = parallel, 90° = perpendicular).

Mechanical forces generated within the cytoskeleton can be transmitted to the nucleus through nucleo-cytoskeletal coupling and can influence nuclear shape and positioning (Lombardi et al., 2011; Maurer & Lammerding, 2019a). To assess the nuclear response to stretch, nuclear orientation and morphology were quantified after 6 hours of cyclic stretch. Nuclear orientation showed the same overall trend as actin filament directionality, with a more concentrated orientation distribution at around 90 degrees in the uniaxially stretched cells compared with the even distributions in unstretched and biaxially stretched cells (Figure 5B). Nuclear shape, measured by aspect ratio, was significantly higher in uniaxially stretched cells compared with unstretched and biaxially stretched cells, indicating nuclear elongation perpendicular to the direction of strain (Figure 5C). However, the nuclear area results showed no significant differences between unstretched, uniaxially stretched, and biaxially stretched cells (Figure 5D).

**Figure 5.**
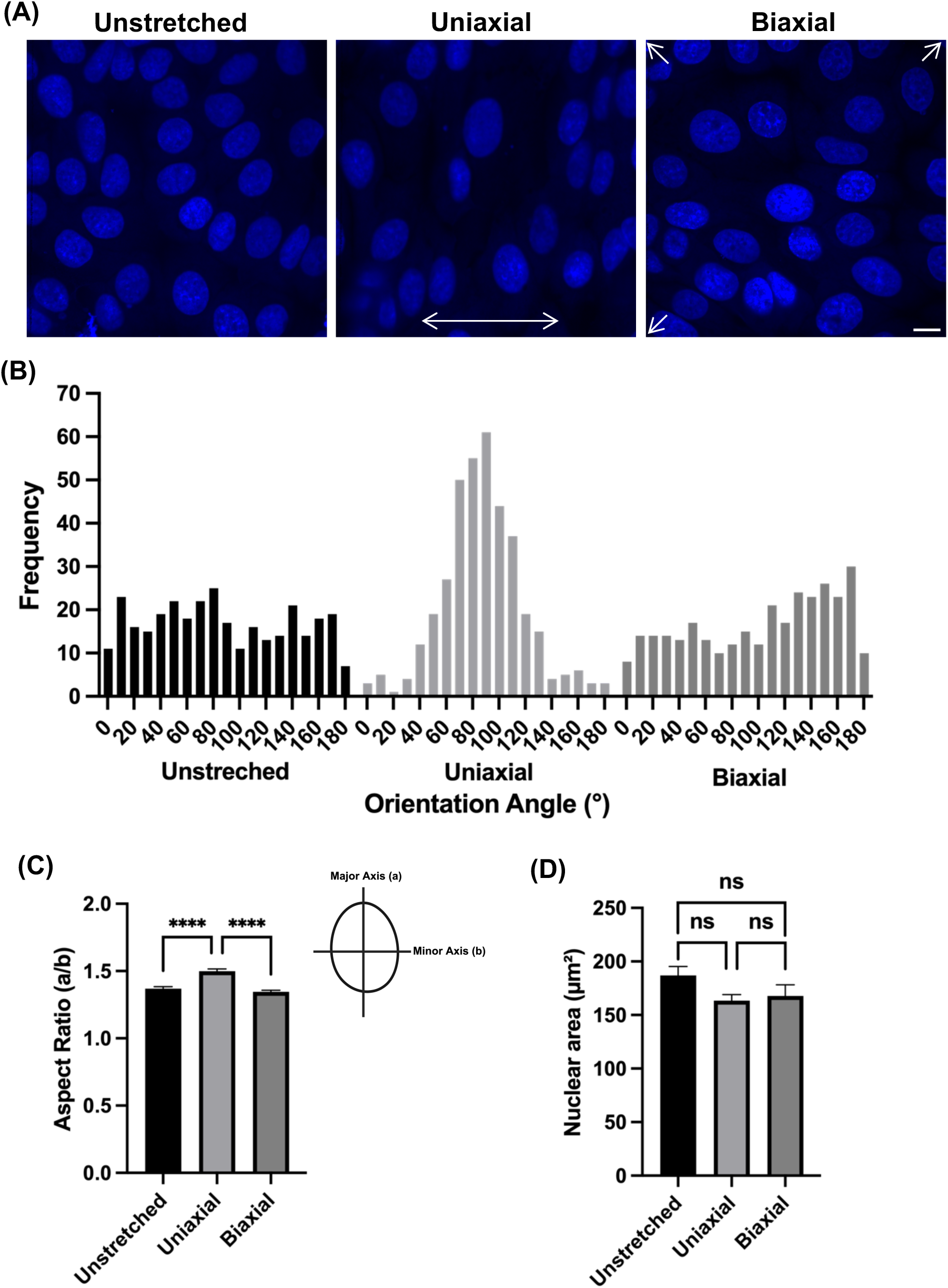
Nuclear orientation and morphology after cyclic stretching of MCF-7 cells. MCF-7 cells were fixed after 6 hours under three conditions: unstretched, uniaxial, and biaxial stretch, then stained with DAPI for nuclear visualization. **(A)** Representative confocal images of DAPI-stained nuclei, scale bar represents 20 μm. **(B)** Nuclear orientation analysis is shown as a frequency distribution of orientation angles. **(C)** Nuclear aspect ratio quantification. **(D)** Nuclear area quantification. Data are presented as mean ± SEM from n = 3 independent experiments. Statistical significance was determined by one-way ANOVA.

## Discussion

This study describes a 3D-printed, motorized, biaxial cell-stretching device designed to apply mechanical strain to adherent cells. Cell-stretching systems are widely used to study mechanotransduction, and cellular responses depend on the direction, magnitude, and type of applied strain (Constantinou & Bastounis, 2023; Kamble et al., 2016a; Wang et al., 2001). Biaxial cell stretching devices expose cells to deformation along two orthogonal axes, applying strain that more closely reflects the mechanical environment of many tissues compared to the single-axis deformation of uniaxial stretching (Gould et al., 2012; Kamble et al., 2016a; Tremblay et al., 2013). Our device builds on the previously developed motorized uniaxial stretcher from our group by using the same motor system and PDMS substrate and adapting the existing MATLAB control program for biaxial stretch (Al-Maslamani et al., 2021). This design allows uniaxial and biaxial stretch responses to be compared using similar experimental setups.

PDMS is widely used in cell-stretching systems because of its elastic and optical properties (Figure 2A) (Constantinou & Bastounis, 2023; Kamble et al., 2016a). Membrane thickness affects how PDMS deforms under load and a consistent well thickness across membranes was needed to ensure the strain applied to cells was comparable between experiments. Our previous membrane casting procedure relied on overflow casting to achieve a thin well bottom. We found the variability between different casts to be too high. Therefore, for the current study, we chose to cast the well walls onto a commercially prepared 250 µm PDMS sheet. Characterization of the membrane showed that deformation increased with motor-arm displacement and was similar along the X and Y axes, confirming symmetrical strain across both axes (Figure 2C).

Having confirmed membrane deformation, we tested whether biaxial stretch produced a cellular response. IEX-1 induction provides biochemical evidence that the device delivers a strain sufficient to drive a transcriptional response. IEX-1 is an immediate-early gene shown to be mechanically regulated by NF-κB in cardiomyocytes under cyclic biaxial strain (De Keulenaer et al., 2002) and to be induced by cyclic strain in vascular smooth muscle cells (Schulze et al., 2003), supporting its use as a mechanosensitive gene. Here, 7% cyclic biaxial stretch increased IEX-1 expression at 1 hour and further at 3 hours, confirming that our device transmitted mechanical strain to the cells.

We next compared how cells responded structurally to uniaxial and biaxial strain. The uniaxial stretcher served as a reference, since cellular responses to cyclic uniaxial stretch are well documented: uniaxial strain reorganizes actin stress fibers and promotes alignment away from the strain axis, toward the direction of minimal substrate deformation (Hayakawa et al., 2001; Hsu et al., 2010; Wang et al., 2001). Our aim was to compare a well-characterized stretching device with the biaxial stretcher under the same experimental conditions. As expected, uniaxial strain produced strong actin alignment perpendicular to the strain direction, consistent with previous reports of stretch-induced reorientation (Figure 4B) (Hayakawa et al., 2001; Hsu et al., 2010; Wang et al., 2001). Biaxially stretched cells showed no dominant actin orientation, but instead showed increased actin signal at the cell periphery and around the nucleus (Figure 4A). This suggests that the actin cytoskeleton response is a more localized remodeling of peripheral and perinuclear actin structures. The stronger peripheral signal may reflect remodeling of the actin cortex, a dynamic actin network that regulates cell shape and resists mechanical stress (Kelkar et al., 2020; Salbreux et al., 2012; Svitkina, 2020). The increased perinuclear actin signal may reflect remodeling of actin structures near the nuclear surface, including the perinuclear actin cap, which connects cytoskeletal organization to nuclear positioning and nuclear shape (Chambliss et al., 2013; Khatau et al., 2010; Kim et al., 2012).

The changes in nuclear shape and orientation under uniaxial strain are consistent with mechanical coupling between the actin cytoskeleton and the nucleus. Forces generated by the cytoskeleton can be transmitted to the nucleus via the LINC complex, enabling extracellular mechanical signals to drive nuclear deformation (Lombardi et al., 2011; Maurer & Lammerding, 2019a). Nuclei in uniaxially stretched cells became more elongated and oriented perpendicular to the axis of strain, similar to the direction of actin reorientation. In contrast, nuclei in biaxially stretched cells showed morphology similar to unstretched controls, consistent with the absence of a dominant actin orientation under biaxial stretching. The nuclear area was not significantly altered under either stretching condition, suggesting that strain affects nuclear shape and orientation without altering nuclear size. These findings support the idea that cytoskeletal organization plays a key role in transmitting mechanical signals to the nucleus (Jaalouk & Lammerding, 2009; Maurer & Lammerding, 2019a).

In conclusion, we have developed a custom 3D-printed, motorized biaxial cell stretching device that delivers biaxial strain to adherent cells *in vitro*. Characterization of the device showed similar deformation levels along the X and Y axes of the membrane. The biological proof-of-concept experiments demonstrated that cyclic biaxial stretching can induce upregulation of the IEX-1 gene in MCF-7 breast cancer cells. Staining and image analyses further showed distinct cytoskeletal and nuclear responses to uniaxial and biaxial stretching, with stronger directional alignment in response to uniaxial stretching than biaxial stretching. This device provides a tool for mechanobiology that allows for a direct comparison of cellular responses to uniaxial and biaxial strain applications. In addition, the design of both stretching devices allow for correlation of microscopy analyses with biochemical data, allowing for a more comprehensive insights into cellular responses.

## Materials And Methods

### Cell Line & Culture

Human breast cancer cell lines MCF-7 (#HTB-22, from ATCC) were cultured in Dulbecco’s Modified Eagle Medium (DMEM) supplemented with 10% (v/v) Fetal Bovine Serum, 1% (v/v) L-Glutamine, and 1% (v/v) penicillin and streptomycin. The cells were maintained at 37°C in a humidified environment containing 5% CO2.

### Biaxial Cell Stretcher

The biaxial stretcher was designed in Autodesk Inventor, and all components were 3D-printed in polylactic acid (PLA; Ultimaker PLA, ø 2.85 mm) using an Ultimaker 3 Extended printer. Printing parameters were: 0.4 mm AA nozzle, 7 mm brim, printing temperature 205 °C, build-plate temperature 60 °C, 100% flow, 2.85 mm filament diameter, print speed 70 mm/s, travel speed 250 mm/s, print acceleration 4000 mm/s², travel acceleration 5000 mm/s², print jerk 25 mm/s, and travel jerk 30 mm/s. Printed parts were left to cool at room temperature overnight. The device was assembled by gluing the four base parts into the main base groove. Four micro linear actuators (Actuonix L12-50-210-6-R) were seated in the motor holders and bolted in place through the motor covers, and motor hands were attached to the actuator arms and secured with screws. The actuator wires were soldered to a circuit board connected to a regulated power supply. The actuators were driven by an Arduino Mega 2560, connected to a computer via USB, and programmed in MATLAB. Cyclic stretch was applied at 7% strain for biaxial and 17% for uniaxial stretching at 0.5 Hz, such that the membrane completed one full stretch-and-return cycle every 2 seconds.

### PDMS Membrane Preparation

PDMS membranes were prepared from DOWSIL 184 silicone elastomer base and curing agent mixed at a 12:1 ratio. The mixture was stirred thoroughly and degassed under vacuum to remove air bubbles. A commercially prepared pre-cured PDMS sheet (HT6240, 40 Shore A, super clear, 0.25 mm thickness) was placed in the membrane mold as the base layer, and the degassed PDMS mixture was poured over it. A square plexiglass plug (20 mm x 20 mm) was placed in the middle of the membranes and the PDMS was poured around this to form the culture well. Membranes were cured at room temperature for 48 h. After curing, membranes were prepared according to stretcher type. For uniaxial stretching, membranes were kept at the mold dimensions and mounting holes were punched. For biaxial stretching, membranes were cut into squares and a hole was punched at each corner for attachment to the four motor arms.

### Fibronectin Coating

PDMS membranes were washed extensively with soap and water, immersed in 70% ethanol for 30 min to sterilize, then transferred to a laminar-flow hood and placed in sterile Petri dishes (150 × 15 mm) to air-dry. For extracellular matrix coating, membranes were incubated with human plasma fibronectin (Gibco, Thermo Fisher Scientific, 33016-015) at 2 µg/mL in phosphate-buffered saline (PBS); 1 mL of fibronectin solution was applied per membrane to cover the culture area. Dishes were sealed with parafilm and stored at 4 °C overnight. Membranes were used for cell seeding the following day.

### Real-Time PCR

MCF-7 cells were seeded at 3.5 × 10⁵ cells per fibronectin-coated membrane and incubated for 48 h at 37 °C and 5% CO₂. Cells were then left unstretched or subjected to 7% biaxial cyclic stretch for 30 min, 1 h, or 3 h, with six independent experiments per time point. Immediately after stretching, cells were washed with PBS, and total RNA was extracted using the RNeasy Plus Mini Kit (Qiagen) according to the manufacturer’s instructions. RNA concentration was measured on a NanoDrop spectrophotometer (Thermo Fisher Scientific), and 1 µg of RNA per sample was reverse-transcribed to cDNA using the High-Capacity cDNA Reverse Transcription Kit (Thermo Fisher Scientific) with the program: 25 °C for 10 min, 37 °C for 120 min, and 85 °C for 5 min.

Real-time PCR was performed on a QuantStudio 6 Flex system using PowerUp SYBR Green Master Mix (Thermo Fisher Scientific, A25742) in 15 µL reactions. Relative expression was calculated by the comparative Ct method, normalized to the housekeeping genes HPRT and PGK1.

Primer sequences were:

IEX-1: forward 5′-AGCCGCAGGGTTCTCTA-3′.

reverse 5′-GATGGTGAGCAGCAGAAAGA-3′.

PGK1: forward 5′-GGAGAACCTCCGCTTTCATGTG-3′.

reverse 5′-GGCTCGGCTTTAACCTTGTTCC-3′.

HPRT: forward 5′-AGGATTTGGAAAGGGTGTTTATTC-3′.

reverse 5′-CCCATCTCCTTCATCACATCTC-3′.

### Actin & DAPI Staining

MCF-7 cells were seeded at 3.5 × 10⁵ cells per fibronectin-coated membrane and incubated for 48 h at 37 °C and 5% CO₂. Cells were then left unstretched, uniaxially stretched at 17%, or biaxially stretched at 7% for 6 hours. Immediately after stretching, cells were washed twice with room-temperature PBS, fixed in 4% paraformaldehyde for 10 min at room temperature, then washed twice with PBS. Cells were permeabilized in PBS containing 0.1% Triton X-100 for 10 min and washed once in PBS. Well bottoms were removed from the PDMS well structure with a scalpel and transferred to a six-well plate containing PBS. For staining, membranes were inverted onto parafilm bearing 100 µL of staining solution (FITC-phalloidin 1:300, DAPI 1:500) and incubated for 1 h. Membranes were washed for 5 min, 4 times, then mounted on slides using ProLong Diamond Antifade Mountant and covered with a coverslip. The slides were kept at room temperature for 1 h and stored at 4 °C overnight. Finally, cells were imaged on a Nikon A1R confocal microscope using VisiView software.

### Image Analyses

Images of stained cells were analyzed using Fiji-ImageJ software. F-actin distribution was measured using the directionality tool, which measures actin filaments based on Fourier spectrum analysis. For nuclear shape and orientation analysis, DAPI-stained images were converted to threshold images, and each nucleus was selected using the wand tool. The ROI manager was used to calculate major and minor axes, orientation, and aspect ratio (a/b).

